# Strong circulation of *Angiostrongylus cantonensis* in a protected Mediterranean wetland. Implications for resident and migratory wildlife

**DOI:** 10.1101/2025.01.30.635808

**Authors:** Marcia Raquel Pegoraro de Macedo, Sebastià Jaume-Ramis, Claudia Paredes-Esquivel

## Abstract

*Angiostrongylus cantonensis*, the rat lungworm, is a metastrongyloid nematode responsible for eosinophilic meningitis and an emerging pathogen of global concern. While its presence has been reported in various regions, including the Canary Islands and mainland Spain, data from the Balearic Islands remain limited. This study investigates the prevalence of *A. cantonensis* in rodents from the Natural Park of s’Albufera, Mallorca, a protected wetland ecosystem. A total of 43 rodents were captured and necropsied, including *Rattus norvegicus, R. rattus* and *Apodemus sylvaticus*. The overall prevalence of *A. cantonensis* in rats was 46.15% (95% CI: 30.43–62.62%), with no significant differences between species, seasons, or sexes. However, adult rats exhibited significantly higher infection rates than juveniles (p = 0.05). Notably, infection intensity and parasite abundance were positively correlated with host weight (ρ = 0.60, p < 0.001). Given the co-occurrence of infected rodents and gastropods, our findings suggest the establishment of *A. cantonensis* in Mallorca, raising concerns about zoonotic transmission risks. The study highlights the need for continued surveillance and public health awareness to mitigate potential human and wildlife exposure in this ecologically significant area.

## 1. Introduction

The rat lungworm, *Angiostrongylus cantonensis*, is a metastrongyloid nematode originally identified infecting rats from Canton, China. It is now recognized as an emerging pathogen in all continents except Antarctica (Cowie *et al*. 2022b; Pandian *et al*. 2023; Jaume-Ramis *et al*. 2023). This parasite is the main etiological agent of eosinophilic meningitis worldwide (Cowie *et al*. 2022a; Wang *et al*. 2008b)

The life cycle of *A. cantonensis* is complex, involving many gastropod species as intermediate hosts and various rat species as definitive hosts. *Rattus rattus* and *R. norvegicus* are the favoured definitive hosts, though at least 17 rodent species may behave as definitive hosts, capable of passing the first stage larvae in their faeces (Barratt *et al*. 2016). Intermediate hosts of *A. cantonensis* include species from as many as 51 mollusc families (Barratt *et al*. 2016; Kim *et al*. 2014). A large number of potential paratenic hosts have also been described (Barratt *et al*. 2016; Cowie *et al*. 2022b). Those play a significant role in the transmission and get infected by consuming intermediate hosts containing L3. Thirty-two paratenic host species including vertebrates and invertebrates (freshwater prawns/shrimp, crayfish, crabs, land planarians, fish, sea snakes, frogs, toads, lizards, centipedes, cattle, pigs and snails) were reported both natural and/or experimentally (Turck *et al*. 2022). The L3 remain infective to definitive and accidental hosts that eat infected paratenic hosts (Barratt *et al*. 2016). Humans get infected, acting as accidental hosts through the ingestion of L3 in raw or undercooked intermediate or paratenic hosts (snails, freshwater shrimps, land crabs, frogs, toads, monitor lizards and planarians), or via vegetables contaminated with gastropods infected with rat lungworm larvae (Hancke *et al*. 2024).

While it was considered a disease of the Far East, reports of locally acquired angiostrongyliasis were spread in sub-tropical and temperate regions and, most recently, *A. cantonensis* was reported in Europe (Galán-Puchades *et al*. 2022; Paredes-Esquivel *et al*. 2019). However, the first report of *A. cantonensis* in the Spanish territory was in the Canary Islands (Foronda *et al*. 2010), followed by Mallorca island, where it was reported in hedgehogs, *Atelerix algirus* (Paredes-Esquivel *et al*. 2019) and gastropods (Jaume-Ramis *et al*. 2023), both naturally infected. Since the first detection in Mallorca, the studies performed have been mainly focused on the detection of the parasite in urban or agricultural areas, following its distribution. In this study we want to identify the presence and prevalence of *A. cantonensis* in a natural protected area, where preliminary rat surveillance has showed a higher prevalence of infection in relation to other parts of the island (unpublished).

## 2. Material and Methods

### 2.1 Ethics statement

The animal protocol and sample collection used in this study was approved and performed under the guidelines set forth by the Institutional Animal Care and Use Committee at the University of the Balearic Islands (CBS-AB 07/2020). Sampling of rats in the Natural Park of s’Albufera was performed under the permission of the Species Protection Service and the Natural Areas Service form the Government of the Balearic Islands (CAP 16/2023).

### 2.2 Sampling

The Natural Park of s’Albufera of Mallorca is located within the municipalities of Muro, Sa Pobla, and Alcudia. It is the largest wetland in the island, with an area of 2,036 hectares, plus a peripheral protection zone and ecological corridors covering areas of 754.30 and 139.03 hectares, respectively (Gobierno 2021).

The rats were sampled in 2023 (November and December) and 2024 (June and July). The park was divided into five sampling zones (FIGURE 1), each approximately 400 hectares in size. In each zone, a total of 90 traps of the Tomahawk (45 × 15 × 17 cm) and Sherman (43 × 12.5 × 14 cm) types were distributed, spaced an average of 50 meters apart, and remained in place for three nights per zone. The sampling effort amounted to 1,350 trap-nights. Peanut butter and *fuet* (a traditional sausage made from pig’s meat) were used together as bait, with daily checks conducted. Animals found in the traps were taken to the University of the Balearic Islands where they were euthanized in a carbon dioxide chamber and frozen at -20°C until the necropsies were performed.

**Figure 1.**
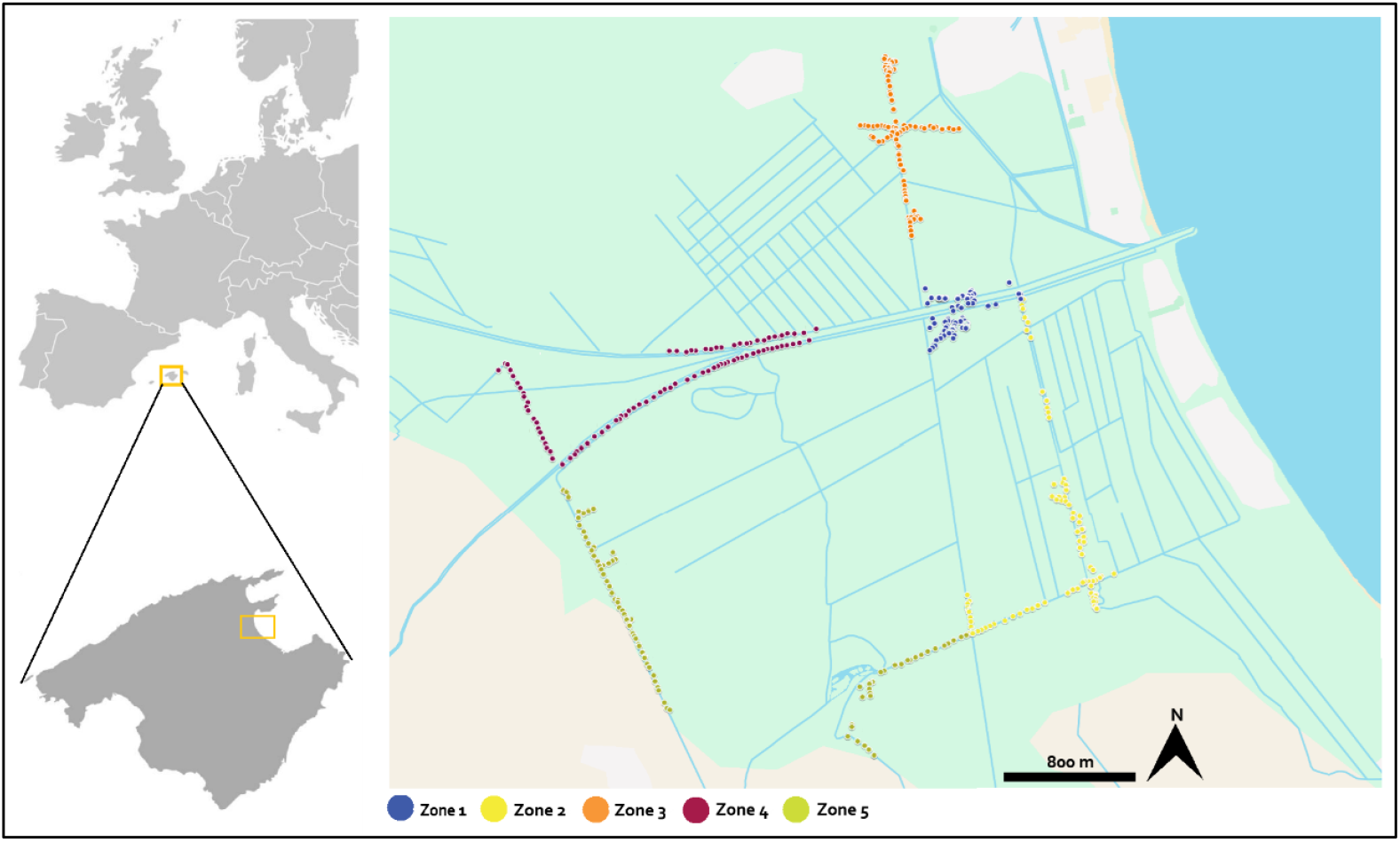
Map of the location of Mallorca Island in relation to Europe; Zoom with the location of the study area, Natural Park of s’Albufera (green area), and the five rodent capture zones.

### 2.3 Necropsy and helminths identitfication

All animals were sexed, weighed and measurements of body length including the nose to anus and the anus to the end of the tail were noted. Species identification was based on morphometric parameters (Bonnefoy *et al*. 2008)

Necropsies were performed in a class II safety cabinet. The heart and lungs were extracted and examined under a stereomicroscope for the presence of parasites. The abdominal and thoracic cavities were also inspected. Nematodes found were identified based on morphological characteristics as described by (Kinsella 1971; Moreira *et al*. 2013) and were preserved in 70% ethanol for subsequent molecular analysis. Morphological identifications were confirmed by molecular analysis, consistent with methods used in previous studies from our laboratory (Delgado-Serra *et al*. 2022)

### 2.4 Statistical analyses

The prevalence of *A. cantonensis* in the rodents was determined for *R. norvegicus* and *R. rattus*, as well as for both species combined. Proportions with 95% confidence intervals were calculated using the function “binom.test” in R v 4.3 (Team 2023). To assess whether there was a statistically significant difference in the prevalence of the *A. cantonensis* among *R. norvegicus* and *R. rattus*, sex, age and season, Fisher’s exact test was employed, using a 2×2 contingency table. The Mann-Whitney U test was used to assess the statistical significance of the difference in infection intensities and average abundance of parasites between *R. norvegicus* and *R. rattus*, sex, age and season. We calculated Spearman’s rank correlation coefficient (rho) to assess the relationship between the weight of and the presence of infection (rho) using the cor.test function in R and between the weight and average abundance of parasites. A variable was considered significant when the p-value was **≤** 0.05.

## 3. Results

We captured and necropsied 43 rodents: 17 *Rattus norvegicus*, 22 *R. rattus*, and 4 *Apodemus sylvaticus*. The overall prevalence of *A. cantonensis* (Figure 2) in the group of rodents was 46.15% (95% CI: 30.43–62.62%) (Table 1). The prevalences for *R. norvegicus* and *R. rattus* were 52.94% (95% CI: 30.96–73.83) and 40.90% (95% CI: 23.25–61.26), respectively. The wild rodent *A. sylvaticus* tested negative for all samples and was not included in the statistical analyses, as to our knowledge, it has not yet been reported as a competent host for *A. cantonensis*.

**Figure 2.**
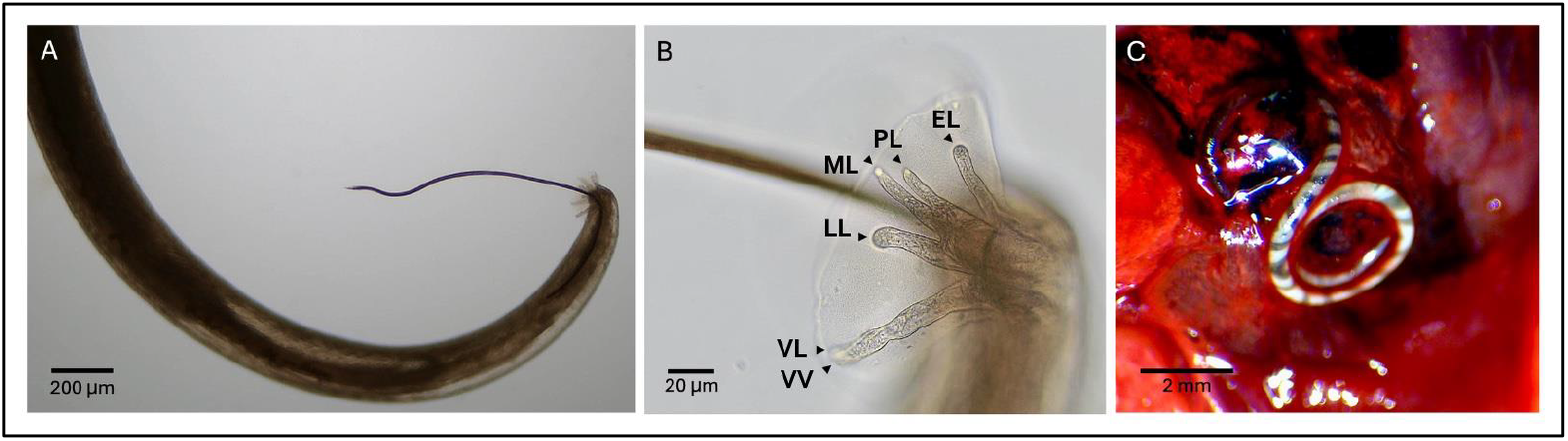
*Angiostrongylus cantonensis*; A. Male, lateral view of caudal bursa spicule. B. Detail of caudal bursa, in lateral view, showing rays disposition: ventroventral (VV), ventrolateral (VL), laterolateral (LL), meiolateral (ML), posterolateral (PL), externodorsal (ED). C. Adult female recovered from lung tissues.

**Table 1.** Prevalence and mean intensity of *A. cantonensis* in rats from Natural Park of s’Albufera, Mallorca, Spain.

| Rats | No. of infected /No. of collected samples | Prevalence % (95% CI) | Mean intensity (worms/host) ± SD |
| --- | --- | --- | --- |
| **Species** |  |  |  |
| Overall | 18/39 | 46.15 (30.43-62.62) | 3.07 ± 4.65 |
| *R. norvegicus* | 9/17 | 52.94 (30.96-73.83) | 5.82 ± 10.20 |
| *R. rattus* | 9/22 | 40.90 (23.25-61.26) | 1.72 ± 4.28 |
| **Sex** |  |  |  |
| Male | 12/18 | 66.67 (41.15-85.64) | 5.35 ± 9.82 |
| Female | 6/20 | 30 (12.83-54.33) | 1.30 ± 3.35 |
| Not identified | 0/1 | - |  |
| **Age** |  |  |  |
| Juvenil | 1/8 | 12.50 (0-52.65) | 0.875 ± 2.48 |
| Adult | 17/31 | 54.83(36.03-72.68) | 4.19 ± 8.36 |
| **Season** |  |  |  |
| Summer | 11/25 | 44 (25.02-64.72) | 4.13 ± 8.77 |
| Winter | 7/14 | 50 (27.4-84.8) | 4.43 ± 5.65 |

Statistical analyses revealed no significant difference in infection rates between *R. norvegicus* and *R. rattus* (p = 0.53), although there was a potential trend toward higher prevalence in *R. rattus* (OR = 1.60; 95% CI: 0.38–7.01). Similarly, no significant differences in prevalence were observed between winter and summer (p = 0.52 95% CI: 0.14–2.70) or between males and females (p = 0.06; OR = 0.26; 95% CI: 0.06–1.08). In terms of age, adults were significantly more likely to be infected than juveniles (p = 0.05), with an approximately eightfold higher likelihood of infection compared to juveniles (OR = 8.10; 95% CI: 0.87–404.18). Although the confidence interval is wide, indicating uncertainty about the exact magnitude of the difference, the statistical significance supports the conclusion that infection rates are significantly higher in adults.

No statistically significant differences were observed in the mean intensity of infection between species (W = 230, p-value = 0.19), age groups (W = 174, p-value = 0.06), or seasons (W = 172, p-value = 0.38). However, males exhibited a significantly higher intensity of infection compared to females (W = 146.5, p-value = 0.04). Similarly, there were no significant differences in the average abundance of parasites when comparing species (W = 230, p-value = 0.19), age groups (W = 74, p-value = 0.06), or seasons (W = 192, p-value = 0.60). Males exhibited a significantly higher average abundance of parasites compared to females (W = 247.5, p-value = 0.034).

A Spearman’s rank correlation analysis revealed a significant association between rat weight and the presence of the parasite (ρ = 0.60, p = 2.449e^-05^), suggesting that heavier rats are more likely to be infected (Figure 3A). Furthermore, a significant positive correlation was observed between the weight of the animals and their parasite abundance (ρ = 0.58, p = 0.0001), suggesting that heavier rats are more likely to have a higher abundance of parasites (Figure 3B).

**Figure 3.**
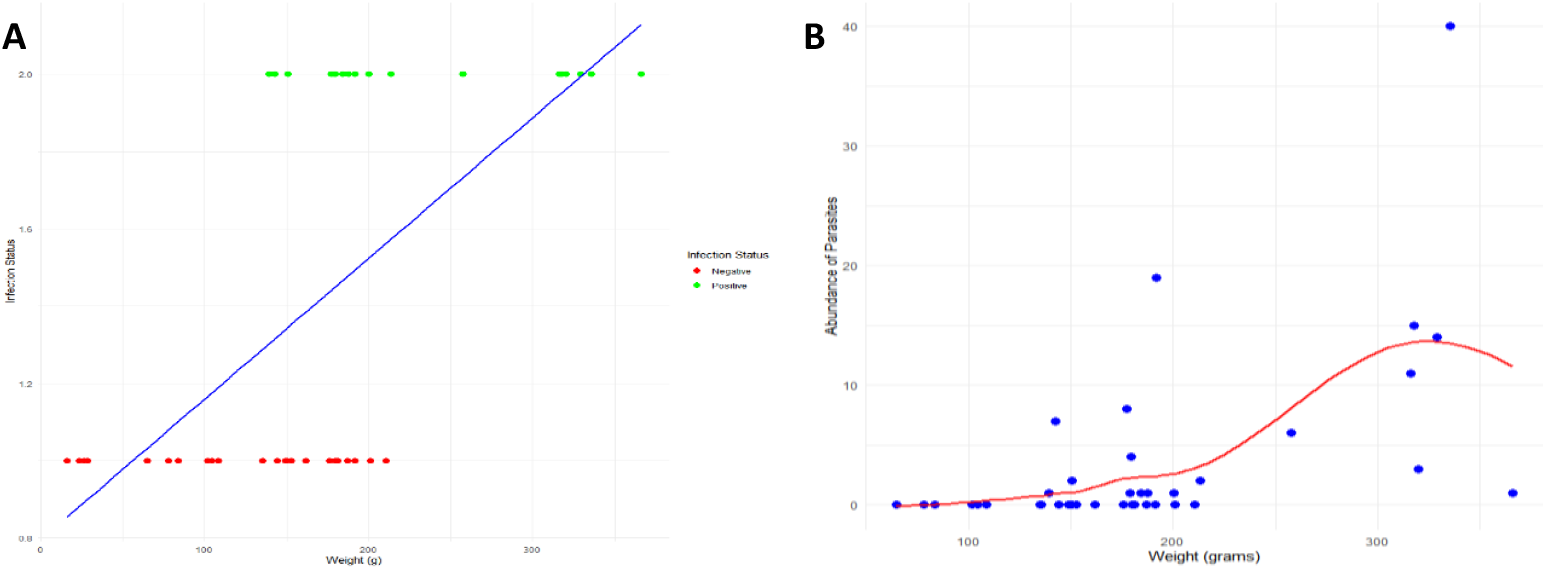
Probability of *Angiostrongylus cantonensis* infection and parasite abundance based on weight. (A) Correlation (blue line) between rat weight and *A. cantonensis* infection status in rats from the Natural Park of s’Albufera, Mallorca. (B) Relationship between rat weight and average parasite abundance, with heavier rats showing higher average abundance. Each dot represents an individual rat.

## 4. Discussion

This survey clearly demonstrates that invasive rats (both *R. norvegicus* and *R. rattus*) in the Natural Park of s’Albufera of Mallorca, act as natural hosts for *A. cantonensis*. Although suggestive of risk, it is not unexpected that the island’s rodents are infected, as infected gastropods have also been reported (Jaume-Ramis *et al*. 2023). Certainly, the co-occurrence of infected rodents and molluscs establishes the ideal scenario for infections in accidental hosts, especially humans, as is commonly observed in the history of neuroangiostrongyliasis (Aguiar *et al*. 1981; Espírito-Santo *et al*. 2013; Liu *et al*. 2024; Pincay *et al*. 2009; Waugh *et al*. 2005).

*Angiostrongylus cantonensis* has been reported in various species of vertebrate and invertebrate hosts since its first description in China in 1935. Continental Europe was considered free of this parasite until 2021, when positive rats were found in Valencia, Spain (Galán-Puchades *et al*. 2022). Spanish island territories have reported *A. cantonensis* for over a decade. In the Canary Islands, located off the coast of North Africa and surrounded by the Atlantic Ocean, *A. cantonensis* has been recorded in humans and animals (Foronda *et al*. 2010; Martin-Alonso *et al*. 2015; Martin-Alonso *et al*. 2011; Martin-Carrillo *et al*. 2023; Martín-Carrillo *et al*. 2021; Segeritz *et al*. 2021). On the island of Mallorca, located in the Mediterranean Sea, hedgehogs of the species *Atelerix algirus* were found with *A. cantonensis* in the central nervous system (Paredes-Esquivel *et al*. 2019) and recently, thirteen species of mollusks were reported as potential intermediate hosts by molecular techniques (Jaume-Ramis *et al*. 2023). Areas where the rat lungworm has been detected in its definitive natural hosts should be considered endemic for *A. cantonensis* (Wang *et al*. 2008a). Infected rats in a conservation area surrounded by agricultural fields and urban landscapes, such as the Natural Park of s’Albufera of Mallorca, are concerning. Given that gastropods are abundant in the park, several risk factors for accidental infection can affect visitors, domestic and wild animals that live there, and residents who work in or consume agricultural products from the nearby area.

It is not well established in the literature when the prevalence of *A. cantonensis* in rats can be considered high, and there appears to be no direct association between rat prevalence and the occurrence of the disease in accidental hosts. In the Philippines, where there were no records of angiostrongyliasis in humans, the prevalence in rodents reached 100% (Castillo, Paller 2018), in Spain it was 20% (Galán-Puchades *et al*. 2022) and in Argentina, 4.3% (Hancke *et al*. 2024). On the other hand, in regions with sporadic cases of angiostrongyliasis in animals and/or humans, rodent prevalence was recorded as 16.5% in Australia (Aghazadeh *et al*. 2015), 22.8% in Florida, USA (Stockdale Walden *et al*. 2017) and 74% in Brazil (Simoes *et al*. 2011).

Our findings offer a new insight into the distribution of rat lungworms in Mallorca and suggest a northward expansion of their range, significantly increasing the risk of disease transmission to humans and wildlife. The overall prevalence of 39.53% with no significant seasonal variation, may be attributed to the abundance and stability of snail populations in the area. The high abundance of mollusks in the park can be linked to its ecological characteristics: the park comprises a shallow freshwater marsh covered by vegetation, an extensive drainage network with hundreds of kilometers of canals, and lagoons. Freshwater sources include both groundwater and surface inflows, with some seawater penetration during the dry season (Mayol 1992). Additionally, the climatic conditions reported by a meteorological station within the park from September 2023 to August 2024—precipitation ranging from 30 to 80 mm in most months and average temperatures between 11°C and 26°C—are conducive to maintaining the mollusk population throughout the year. Favorable conditions for the occurrence of eosinophilic meningoecephalitis caused by *A. cantonensis* align with the parasite’s life cycle and focus on the living conditions of the intermediate host (Dorta-Contreras *et al*. 2015). In Cuba, despite the lack of a tradition of consuming mollusks, higher concentrations of infections were recorded in years when precipitation ranged from 30 to 170 mm and temperatures ranged from 22°C to 28°C (Dorta-Contreras *et al*. 2015). Given the favourable meteorological conditions in the region, it is crucial to remain vigilant about the risk of infection year-round.

Rodents are indispensable for the establishment of *A. cantonensis* foci in a given region, as they serve as a persistent source of infection, sustaining the parasite’s life cycle within the area. *A. cantonensis* has evolved into a major emerging pathogen globally, driven by recurrent outbreaks and a rising number of sporadic case reports, autochthonous and travel-related, in previously unaffected regions. However, there have been no reported cases of angiostrongyliasis in Mallorca so far. The study and the vigilance of the life cycle of *A. cantonensis* in the island must be established first in order to understand how to control and prevent angiostrongyliasis.

The detection of the rat lungworm in the region raises concerns beyond the potential risk of angiostrongyliasis in humans. Four other rodent species, including wild species, coexist on the island: *Apodemus sylvaticus* (Linnaeus, 1758), *Eliomys quercinus* (Linnaeus, 1766), *Mus musculus* (Linnaeus, 1758), and *Mus spretus* (Lataste, 1883)(Palomo *et al*. 2007). *A. sylvaticus* was negative in our study but is a natural definitive host of *Angiostrongylus dujardini* (Alvarez *et al*. 1991; Eira *et al*. 2006). *Mus musculus* has been recorded as an incidental host of *A. cantonensis* in the Canary Islands (Martin-Carrillo *et al*. 2023), but it is currently considered a competent host in experimental infections. *Mus spretus* is not recorded as a host of *A. cantonensis*, but it has been reported to harbor *Angiostrongylus* sp. in the southeastern Iberian Peninsula (Torres *et al*. 2003). *Eliomys quercinus* has not yet been documented as a host for *Angiostrongylus*. In addition to rodents, other mammals that consume mollusks and are present on Mallorca Island include *Genetta genetta* (Linnaeus, 1758) and *Lepus granatensis* (Rosenhauer, 1856) (Palomo *et al*. 2007). Consequently, veterinary professionals and institutions are advised to pay close attention to and report any clinical signs of neuroangiostrongyliasis should they occur in these species.

Considering the history of indigenous angiostrongyliasis and the occurrence of positive rodents near human or animal residences (Espírito-Santo *et al*. 2013; Rivory *et al*. 2024; Rivory *et al*. 2023; Simoes *et al*. 2011; Stockdale Walden *et al*. 2017) it is pertinent to highlight the finding of two positive animals in capture stations very close to agricultural fields and residences. The movement of A. cantonensis to new regions is primarily mediated by male rats, which often have greater dispersal ability than females, while females are important for maintaining *A. cantonensis* on a small, local scale (Simões *et al*. 2014). Additionally, the Natural Resources Management Plan (Gobierno 2021) provides for the use of organic sludge from dredging for agricultural fertilization, a practice that could transport larvae or positive mollusks from the park area to agricultural zones on the island, facilitating the spread of *A. cantonensis*.

## 5. Conclusion

The distribution of *A. cantonensis* in Spain is undeniably facilitated by globalization, climate change and environmental management. Based on our findings, we emphasize the need for health education programs, proper environmental management, attention to diagnosis, and continuous epidemiological surveillance. The adoption of good agricultural practices and awareness of the safe consumption of molluscs and food hygiene will be crucial for preventing the transmission of angiostrongyliasis in Mallorca. Additionally, epidemiological studies on rodent populations in Europe, particularly in urban and peri-urban areas, along with the implementation of effective rodent control measures, are essential to minimize the risk of future infections.

## Acknowledgments

We would like to thank Dr. Guillem Xavier Pons of the *Societat d’Història Natural de les Balears* for providing the Sherman traps used in this study. Thanks to the members of the COFIB for its assistance on the sampling and to Pere Vicens, naturalist of the Natural Park of s’Albufera of Mallorca, for all his help during the study, as well to all the staff of the park.

